# Targeting Fatty Acid Desaturase I Inhibits Renal Cancer Growth Via ATF3-mediated ER Stress Response

**DOI:** 10.1101/2024.03.23.586426

**Authors:** Gioia Heravi, Zhenjie Liu, Mackenzie Herroon, Alexis Wilson, Yang-Yi Fan, Yang Jiang, Nivisa Vakeesan, Li Tao, Zheyun Peng, Kezhong Zhang, Jing Li, Robert S. Chapkin, Izabela Podgorski, Wanqing Liu

**Affiliations:** Department of Pharmaceutical Sciences, Eugene Applebaum College of Pharmacy and Health Sciences, Wayne State University, Detroit, MI 48201, USA; Department of Pharmacology, School of Medicine, Wayne State University, Detroit, MI 48201, USA; Department of Nutrition, Program in Integrative Nutrition and Complex Diseases, Texas A&M University, College Station, TX, 77843, USA; Department of Physiology, Wayne State University School of Medicine, Detroit, MI 48201, USA; Center for Molecular Medicine and Genetics, Wayne State University, Detroit, MI 48201, USA; Department of Biochemistry, Microbiology, and Immunology, School of Medicine, Wayne State University, Detroit, MI 48201, USA; Department of Oncology, School of Medicine, Wayne State University, and Karmanos Cancer Institute, Detroit, MI 48201, USA; CPRIT Regional Center of Excellence in Cancer Research, Texas A&M University, College Station, TX, 77843, USA

**Keywords:** FADS1, PUFA, kidney cancer, ER stress, ATF3

## Abstract

Monounsaturated fatty acids (MUFAs) play a pivotal role in maintaining endoplasmic reticulum (ER) homeostasis, an emerging hallmark of cancer. However, the role of polyunsaturated fatty acid (PUFAs) desaturation in persistent ER stress driven by oncogenic abnormalities remains elusive. Fatty Acid Desaturase 1 (FADS1) is a rate-limiting enzyme controlling the bioproduction of long-chain PUFAs. Our previous research has demonstrated the significant role of FADS1 in cancer survival, especially in kidney cancers. We explored the underlying mechanism in this study. We found that pharmacological inhibition or knockdown of the expression of FADS1 effectively inhibits renal cancer cell proliferation and induces cell cycle arrest. The stable knockdown of FADS1 also significantly inhibits tumor formation *in vivo*. Mechanistically, we show that while FADS1 inhibition induces ER stress, its expression is also augmented by ER-stress inducers. Notably, FADS1-inhibition sensitized cellular response to ER stress inducers, providing evidence of FADS1’s role in modulating the ER stress response in cancer cells. We show that, while FADS1 inhibition-induced ER stress leads to activation of ATF3, ATF3-knockdown rescues the FADS1 inhibition-induced ER stress and cell growth suppression. In addition, FADS1 inhibition results in the impaired biosynthesis of nucleotides and decreases the level of UPD-N-Acetylglucosamine, a critical mediator of the unfolded protein response. Our findings suggest that PUFA desaturation is crucial for rescuing cancer cells from persistent ER stress, supporting FADS1 as a new therapeutic target.

## Introduction

Clear cell renal cell carcinoma (ccRCC) is the most common subtype of kidney cancer. The 5-year survival rate for ccRCC is 50-69%, and for distant stages the survival rate drops to 10% (1). Treatment for localized RCC involves surgery (2), but about 30% of patients eventually develop metastatic RCC (mRCC), which is resistant to conventional chemotherapy (3). The current first-line treatment for mRCC is a combination of immunotherapy and anti-angiogenic tyrosine kinase inhibitors targeting vascular endothelial growth factor (VEGF) (4–6). However, these targeted therapies have limitations and the need for new treatment modalities is evident.

Metabolic reprogramming, a hallmark of most types of cancer, plays a vital role in cancer cell growth, energy metabolism, storage, and signaling (7–9). Targeting genes or enzymes involved in metabolism could be a promising strategy to inhibit cancer cell growth (10,11). Previous studies have demonstrated that fatty acid metabolism, especially the desaturation process, is critical for cancer progression and therapy evasion (12,13). This is consistent with the role of Stearoyl-CoA Desaturase (SCD), the rate-limiting enzyme that catalyzes the conversion of saturated fatty acids (SFAs) into monounsaturated fatty acids (MUFAs) (14,15). Several studies have shown that SCD plays a role in tumor progression via modulation of endoplasmic reticulum (ER) homeostasis (16–19). Interestingly, supplementing polyunsaturated fatty acids (PUFAs) *in vitro* attenuated the ER stress caused by SCD inhibition (20). Similarly, a decrease in long-chain PUFAs (LC-PUFAs) production disrupts ER homeostasis (21). PUFAs, especially LC-PUFAs such as arachidonic acid (AA), eicosatetraenoic acid (EPA), and docosahexaenoic acid (DHA), are important signaling molecules and key components of biomembrane phospholipids (22,23). PUFAs and their derivatives have been implicated to play broad functions in cancer impacting various stages from tumorigenesis to progression and metastasis (24–28). Previous research has shown that Fatty Acid Desaturase 1 (FADS1), a rate-limiting enzyme in LC-PUFA bioproduction, plays a significant role in the growth of various cancer cells (13) and is overexpressed in colon, pancreas, breast, and laryngeal cancers (29). However, whether the LC-PUFA production, controlled by FADS1, plays a similar role as MUFAs remains to be explored. Recently, we showed that FADS1 gene expression is associated with poor patient survival in multiple cancer types, and especially in kidney cancers (13). Moreover, we have also demonstrated that the FADS1 inhibitor (D5D-IN-326) was able to reduce the cell proliferation in ccRCC cell lines (13).

In this study, we further investigated the role and downstream mechanism of action of FADS1 on renal cancer cell proliferation and tumorigenesis. Our findings demonstrate that inhibiting FADS1 activity or knockdown of FADS1 expression can impede renal cancer cell proliferation and induce cell cycle arrest. Our *in vivo* mouse tumorigenesis experiments further indicate that reduced FADS1 expression reduces tumor formation and growth. Mechanistically, we show that FADS1 downregulation results in inhibition of tumor growth through induction of ATF3-mediated ER stress response. Collectively, our findings suggest that FADS1 could be a novel therapeutic target for renal cancer carcinoma.

## Methods

### Cell culture

The human renal cancer cell lines (786-o, A498, ACHN, Caki-1), RPTEC, and HEK 293 were purchased from the American Type Culture Collection (ATCC, VA, USA). 786-o, A498, ACHN, and HEK 293 were maintained in Dulbecco’s modified Eagle’s medium (DMEM, Sigma-Aldrich, St Louis, MO, USA) supplemented with 10% fetal bovine serum (FBS, Thermo Fisher Scientific, Waltham, MA, USA) and 1% penicillin-streptomycin solution (Life Technologies, Carlsbad, CA, USA). Caki-1 was maintained in McCoy’s 5A medium (Life Technologies, Carlsbad, CA, USA) with 10% FBS and 1% penicillin-streptomycin solution. RPTEC was maintained in basal medium (ATCC, VA, USA) consisting of renal epithelial cell growth kit (ATCC, VA, USA) and 1% penicillin-streptomycin solution. All cells were cultured in a 5% CO_2_ 37 °C humidified incubator.

### Transfection

786-o and A498 cells were seeded in 12-well plates (Fisher Scientific, Hampton, NH, USA) and transfected when 70% confluent. 1x 10^5^ infectious units of specific shRNA lentivirus particles (sh-scramble, sh-FADS1, or sh-ATF3; Santa Cruz Biotechnology, Dallas, TX, USA; detail sequence information of each shRNA was shown in the Supplemental Table S1) were added into the culture medium with 5 μg/mL polybrene (Santa Cruz Biotechnology, Dallas, TX, USA). Transfection was discontinued by changing the culture medium 24 hours post treatment. Cells were treated with 10 μg/mL puromycin dihydrochloride (Santa Cruz Biotechnology, Dallas, TX, USA) for 24 hours to elute the non-transfected cells. The condition of the gene and protein knockdown were evaluated by qPCR assay and western blot assay which is shown in the “RNA extraction and RT-qPCR” and “Western blot” section.

### FADS1 inhibition assay

To investigate the impact of FADS1 in cancer cell growth, equal amounts of renal cancer cells, including the transfected 786-o and A498 cells, were cultured with increasing concentrations (20, 200, 2,000 nM; Dimethyl sulfoxide [DMSO] as vehicle) of D5D-IN-326 (FADS1 inhibitor; Sigma-Aldrich, St Louis, MO, USA) for 96 hours. The total cell number and condition of the cells were counted with hemocytometer or evaluated by the immunofluorescence assay which was showed in the “Immunofluorescence staining” section.

### ER stress stimulation assay

To investigate the role of ER stress-induced FADS1 expression in cancer cell growth, the 786-o and A498 cells were treated with increasing concentrations of ER stress stimulators separately; 1 ng/mL or 10 ng/mL Tunicamycin (TM; Cell Signaling Technology, Danvers, MA, USA), 1 nM or 10 nM Thapsigargin (TG; Cell Signaling Technology, Danvers, MA, USA), DMSO as vehicle for both. The qPCR assay was used to evaluate the ER stress associated genes and FADS1 gene expression, which was shown in the “RNA extraction and RT-qPCR” section.

To investigate the function of FADS1 in ER stress-induced cancer cell apoptosis and reduced growth, 786-o and A498 cells were treated with the low dose ER stress agents separately (1 ng/mL TM or 1 nM TG) for 18 h. In some experiments, 2,000 nM D5D-IN-326 (DMSO as vehicle) was co-treated with the ER stress induced cells for an additional 48 h. Cell apoptosis and proliferation were evaluated by the immunofluorescence assay, as described in the “Immunofluorescence staining” section.

### Cell cycle assay

Short hairpin RNA (shRNA) (sh-scramble or sh-FADS1) or D5D-IN-326 (DMSO as vehicle) treated 786-o and A498 cells were cultured in a 12-well plate for 96 hours. The cells were washed in a cold PBS solution (Thermo Fisher Scientific, Waltham, MA, USA), followed with propidium iodide (PI, Abcam biotechnology company, Cambridge, UK) containing RNase staining for 20 minutes at 37 °C. Flow cytometry (Microscopy, Imaging & Cytometry Resources core facility at Karmanos Cancer Institute, Wayne State University) was used to detect the PI staining intensity, which was used to evaluate the cell cycle distribution.

### RNA extraction and RT-qPCR

The 786-o or A498 cells treated with shRNA (sh-scramble or sh-FADS1), D5D-IN-326 (DMSO as vehicle), or ER stress simulator were cultured in a 6-well plate for 48 h. RNA was extracted using the RNA mini kit (Qiagen, Hilden, Germany). cDNA was synthesized using the reverse transcription kit (Qiagen, Hilden, Germany) and quantitative RT-PCR was performed using gene-specific primers (see Supplemental Table S2 for details) and SYBR Green master mix (Thermo Fisher Scientific, Waltham, MA, USA). Negative delta cycle threshold (-Δ Ct) values were calculated by using the housekeeping gene (*PPIA*).

### Immunofluorescence staining

ShRNA (sh-scramble, sh-ATF3, or sh-FADS1), D5D-IN-326 (DMSO as vehicle), or ER stress simulator treated 786-o and A498 cells were cultured on glass coverslips for 96 h. Cells were fixed with 4% paraformaldehyde (Fisher Scientific, Hampton, NH, USA) for 10 min at room temperature. Following PBS washing, fixed cells were incubated with PBS solution containing 5% donkey serum (Jackson ImmunoResearch Labs, West Grove, PA, USA) and 0.2% Triton X-100 (Thermo Fisher Scientific, Waltham, MA, USA) for 1 h at room temperature. Cells were then incubated with primary antibodies overnight at 4°C (the primary antibodies are listed in Supplemental Table S3). The following day, cells were incubated with secondary antibodies for 2 h at room temperature (the secondary antibodies are listed in the Supplemental Table S4). 4′,6-diamidino-2-phenylindole (DAPI; Thermo Fisher Scientific, Waltham, MA, USA) was used to label all nuclei. All samples were observed and imaged by Keyence microscope (Keyence, IL, USA). Low magnification images were used to check the total condition of the cells in each well, while high magnification images were used to assess protein expression.

### Western blot

ShRNA (sh-scramble or sh-FADS1), D5D-IN-326 (DMSO as vehicle), or ER stress stimulator treated 786-o and A498 cells were cultured in a 6-well plate for 48 h. Cells were lysed in RIPA buffer (Thermo Fisher Scientific, Waltham, MA, USA) which contained a complete protease inhibitor cocktail for 30 min at 4 °C. The resulting lysates were centrifuged at 12,000 g for 10 min and supernatants were collected. The protein concentration of each lysate was determined by the Pierce BCA protein assay kit (Thermo Fisher Scientific, Waltham, MA, USA). Cell lysates (20 μg) were analyzed via electrophoresis using 4%-12% sodium dodecyl sulphate-polyacrylamide gradient gels (Thermo Fisher Scientific, Waltham, MA, USA). Pre-stained protein ladders (Thermo Fisher Scientific, Waltham, MA, USA) were used to determine the protein size. Proteins were transferred to nitrocellulose membranes (Thermo Fisher Scientific, Waltham, MA, USA) and blocked with 5% non-fat milk (Thermo Fisher Scientific, Waltham, MA, USA) in TBST (Thermo Fisher Scientific, Waltham, MA, USA) for 30 min at room temperature. Different primary antibodies were incubated with membranes over night at 4 °C (the primary antibodies are listed in the Supplemental Table S5). Following TBST washing, membranes were incubated with appropriate secondary antibodies for 1 h at room temperature (the secondary antibodies are listed in Supplemental Table S6). The ChemiGlow West Chemiluminescence Substrate kit (Thermo Fisher Scientific, Waltham, MA, USA) was used to detect chemiluminescent signals using a ChemiDoc imaging system. ImageJ (NIH, USA) was used to quantify protein bands on blot images. Housekeeping proteins (GAPDH or Vinculin) were used as an internal control to normalize the protein expression data.

### Animal study

*In vivo* xenograft studies were performed on 7-8-weeks-old male and female SCID mice purchased from Charles River laboratories (Wilmington, MA). All experiments involving mice were performed in accordance with the protocol approved by the institutional Animal Investigational Committee of Wayne State University and NIH guidelines (IACUC-21-12-4269). 786-o cells were injected subcutaneously into both the right and left flanks of the mice, with the same mouse receiving both sh-scramble and sh-FADS1 cells to serve as an internal control. Cells in PBS were mixed 1:1 with Cultrex ^TM^ (R&D systems, Minneapolis, MN, USA) at a final concentration of 1 x 10^6^ in 100 μl prior to injection. Control (WT) cells were injected in the right flank, KO (sh-FADS1) were injected in the left flank. Two weeks post injection, tumors were palpated and, if measurable, measured via caliper twice weekly. After 8 weeks, the mice were euthanized, the tumors were resected and measured (weight and size), imaged, and either snap frozen in liquid nitrogen for RNA analysis, or fixed in Z-fix (Anatech, Battle Creek, MI, USA) for embedding and histology.

### Lipidomic analysis

786-o sh-scramble and sh-FADS1 cells 786-o cells were treated with 2 µM D5D-IN-326 (DMSO as vehicle) for 48 h. Subsequently, total phospholipids were separated on silica gel plates using 90 : 8 : 1 : 0.8 (volume ratio) chloroform / methanol / acetic acid / water. The isolated total phospholipids were transesterified using 6% methanolic hydrochloric acid over night at 80 °C. Gas chromatography-mass spectrometry was used to analyze the ratio of arachidonic acid/dihomo-γ-linolenic acid (AA/DGLA) according to previously established methods (30).

### Metabolomic analysis

786-o sh-scramble and sh-FADS1 cells were seeded and cultured to 90% confluence. The cells were rapidly (<15 s) rinsed twice with warm PBS to remove the culture medium, ensuring minimal leakage of intracellular metabolites. Subsequently, the entire cell culture dish was immersed in liquid nitrogen to quickly halt metabolic activity. Metabolites in frozen cells were extracted twice with 80% methanol (pre-cooled at −80°C). Supernatant from two extractions were combined and dried in a CentriVap® vacuum concentrator (Kansas City, MO) at 6°C. The dried cell extract was reconstituted and subject to LC-MS/MS based targeted metabolomic profiling of ∼ 250 water soluble metabolites in the Pharmacology and Metabolomics Core at Karmanos Cancer Institute, as described previously (31). Metabolite concentrations were normalized to protein concentration of each cell sample. Data analysis was carried out using the MetaboAnalyst software (https://www.metaboanalyst.ca/).

### Quantification study and statistical analysis

All biological samples were collected from at least 3-6 independent experiments. The immunofluorescence intensity of nuclei in each well was analyzed by ImageJ software (NIH, USA). DAPI, Ki-67, and CASP positive cells were counted using the cell counter plugin of the ImageJ software. The percentage of positive cells were calculated as (number of the positive cells) * 100% / (number of the DAPI positive cells). For RT-qPCR, negative delta cycle threshold (-Δ Ct) values were calculated by using the housekeeping gene (*PPIA*). The -Δ Ct values were normalized by calculating the z-scores of each gene to create the heatmaps. The 2^(- ΔΔ Ct) was used to determine the relative quantification of gene expression. For Western blots, ImageJ was used to quantify protein bands on each blot image. Housekeeping proteins (GAPDH or Vinculin) was used as an internal control to normalize the protein expression. For flow cytometry, flowJo software (BD, Franklin Lakes, NJ, USA) was used to quantify cell populations. The Dean-Jett-Fox algorithm was used to model the cell cycle and calculate the proportion of each phase. For statistical analysis, data were analyzed using Graphpad Prism software (Boston, MA, USA), which included two tails, unpaired student’s t test or one way ANOVA test followed with Tukey comparison. P-values less than 0.05 were considered statistically significant. For comparisons, data were expressed as fold change relative to the control group. The figure’s diagram was generated using BioRender (Science Suite Inc, Toronto, ON, Canada).

## Results

### Inhibiting FADS1 activity suppresses renal cancer cell proliferation in vitro

In this study, we utilized a recently developed small molecule FADS1 inhibitor D5D-IN-326 to inhibit FADS1 activity. This inhibitor has demonstrated selectivity for FADS1 in previous studies (13). Treatment with D5D-IN-326 significantly inhibits the conversion of dihomo-γ-linolenic acid (DGLA), the direct substrate of FADS1, into arachidonic acid (AA), as indicated by the reduced AA/DLGA ratio in the renal cancer cell line 786-o (**Supplementary Fig. S1**). To investigate the impact of FADS1 inhibition on renal cancer cell growth, we treated human embryonic kidney cell line (HEK293), human renal proximal tubule epithelial cells (RPTEC), and various renal cancer cell lines (786-o, A498, ACHN, and Caki-1) with increasing concentrations of D5D-IN-326 for 96 h (**Fig. 1A**). The results show that D5D-IN-326 treatment significantly reduced the number of renal cancer cells, while it has minimal effect on normal RPTEC renal cells or HEK293 cells (**Fig. 1B**). We chose 786-o and A498 cells for further evaluation, since both are classified as typical primary ccRCC cells (32).

**Figure 1.**
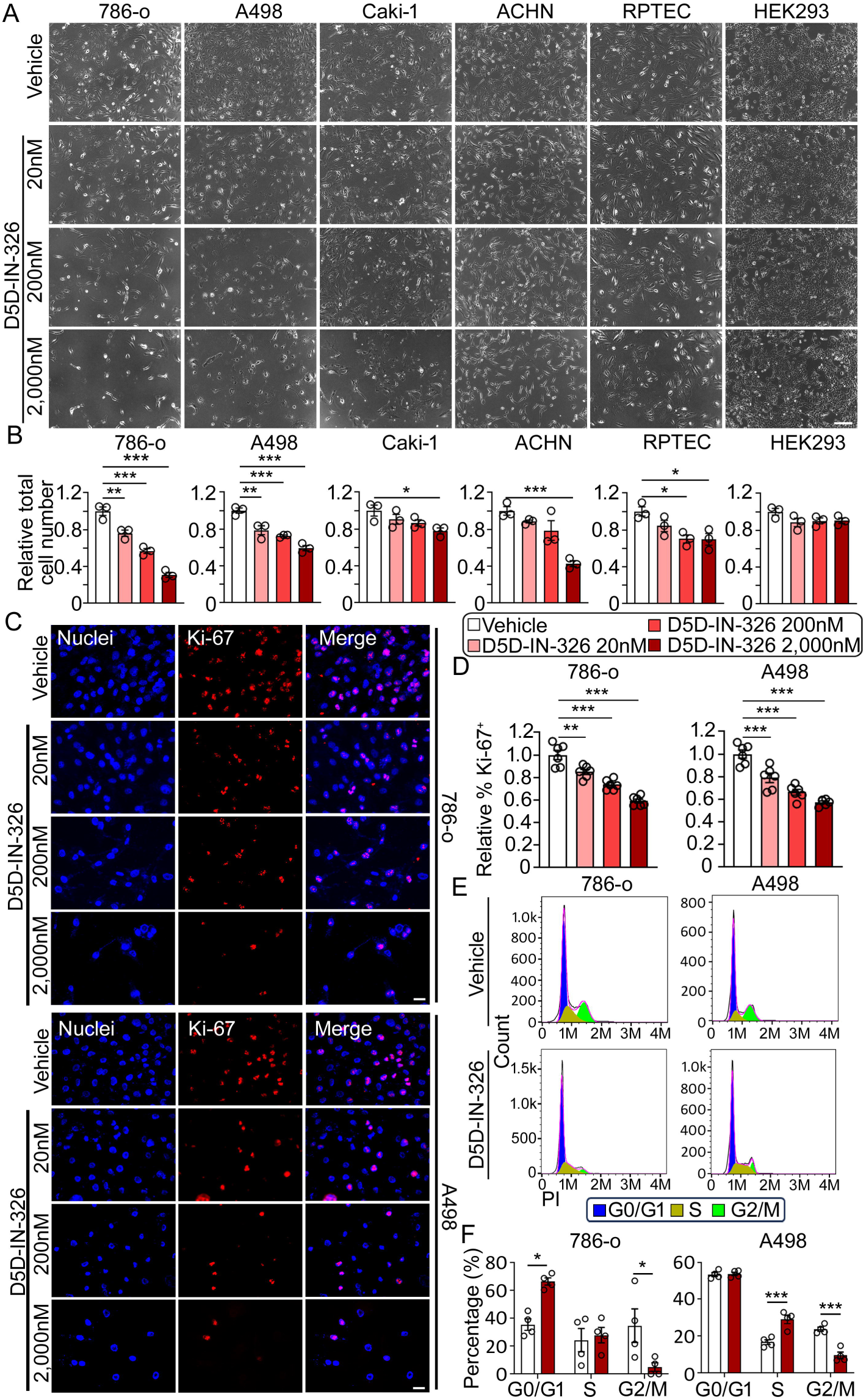
Inhibiting Fatty Acid Desaturase 1 (FADS1) suppressed human renal cancer cell growth *in vitro*. (A) Representative phase images of human renal cancer cells (786-o, A498, Caki-1, and ACHN) and human normal cells (RPTEC and HEK293) treated with increasing concentrations (vehicle, 20, 200, and 2,000 nM) of D5D-IN-326 (FADS1 inhibitor) for 96 hours. Scale bar: 200 μm. (B) The column bar graphs showing the relative quantification of total cell numbers following D5D-IN-326 (FADS1 inhibitor) treatment for 96 hours (mean ± standard error, data were normalized to the vehicle group). Statistical analysis was conducted using Tukey’s multiple comparisons test. *P<0.05; **P<0.01; ***P<0.001. (C) Representative immunofluorescence images illustrating Ki-67 expression in human renal cancer cells (786-o; top panels & A498; bottom panels) treated with increasing concentrations (vehicle, 20, 200, and 2,000 nM) of D5D-IN-326 for 96 hours. Scale bar: 10 μm. (D) The column bar graphs showing the relative quantification of the percentage of Ki-67 positive cells in human renal cancer cells (786-o; left & A498; right) subjected to gradient D5D-IN-326 treatment for 96 hours (mean ± standard error). Data were shown as normalized ratios to the vehicle group. Statistical analysis conducted using Tukey’s multiple comparisons test. **P<0.01; ***P<0.001. (E) Representative histogram showing the expression of the propidium iodide (PI) in human renal cancer cells (786-o; left & A498; right) subjected to gradient D5D-IN-326 treatment for 96 hours. The different stages of cell cycle (G0/G1, S, and G2/M stage) were determined by the expression of PI, as shown in the plot. (F) The column bar graphs showing the percentage of cells (786-o; left & A498; right) distributed in each cell cycle stage (G0/G1, S, and G2/M) after the D5D-IN-326 treatment for 96 hours (mean ± standard error). Statistical analysis conducted using two-tailed unpaired Student’s t test. *P<0.05; ***P<0.001.

To investigate whether FADS1 inhibition affects tumor cell proliferation, we performed immunofluorescence staining of proliferation marker Ki-67 in 786-o and A498 cells treated with increasing concentrations of the FADS1 inhibitor for 96 h. The data revealed a significant, dose-dependent decrease in the percentage of Ki-67 positive (Ki-67^+^) cells upon FADS1 inhibitor treatment compared to vehicle control (**Fig. 1C & 1D**). Additionally, cell cycle analysis showed that a significant fraction of 786-o cells treated with the FADS1 inhibitor (2,000 nM) was arrested in the G0/G1 stage, while fewer cells were arrested in the G2/M stage. In addition, A498 cells treated with the FADS1 inhibitor (2,000 nM) were arrested in the S stage, with fewer cells arrested in the G2/M stage (**Fig. 1E & 1F**). Collectively, these results indicate that FADS1 inhibition reduces renal cancer cell proliferation by inducing cell cycle arrest at the interphase stages preparing cell growth and DNA synthesis prior to cell mitosis.

### Renal cancer cell proliferation is inhibited by stable FADS1 knockdown

To further verify the impact of FADS1 inhibition on RCC cell proliferation and to elucidate the role of FADS1 in renal cancer cell growth, we employed FADS1-specific shRNA to knock down FADS1 expression in 786-o and A498 cells. FADS1 mRNA and protein expression in sh-FADS1 786-o and A498 cells were significantly reduced compared to the cells treated with scrambled shRNAs (**Supplementary Fig. S2 & S3**). As expected, FADS1-knockdown (FADS1-KD or sh-FADS1) significantly reduced the ratio of AA/DGLA in the renal cancer cell line 786-o (**Supplementary Fig. S4**). Consequently, the total number of FADS1-KD 786-o and A498 cells was significantly lower than the scramble shRNA-edited cells, and the percentage of Ki-67^+^ cells was also significantly reduced (**Fig. 2A, 2B, & 2C**). Moreover, cell cycle analysis revealed that FADS1 knockdown led to cell cycle arrest at the G0/G1 stage in 786-o cells and at the S stage in A498 cells (**Fig. 2D & 2E**), mirroring the effects observed with FADS1 inhibitor treatment. These findings confirm the growth suppressing effects of pharmacological FADS1 inhibition and underscore the crucial role of FADS1 gene expression in renal cancer cell growth.

**Figure 2.**
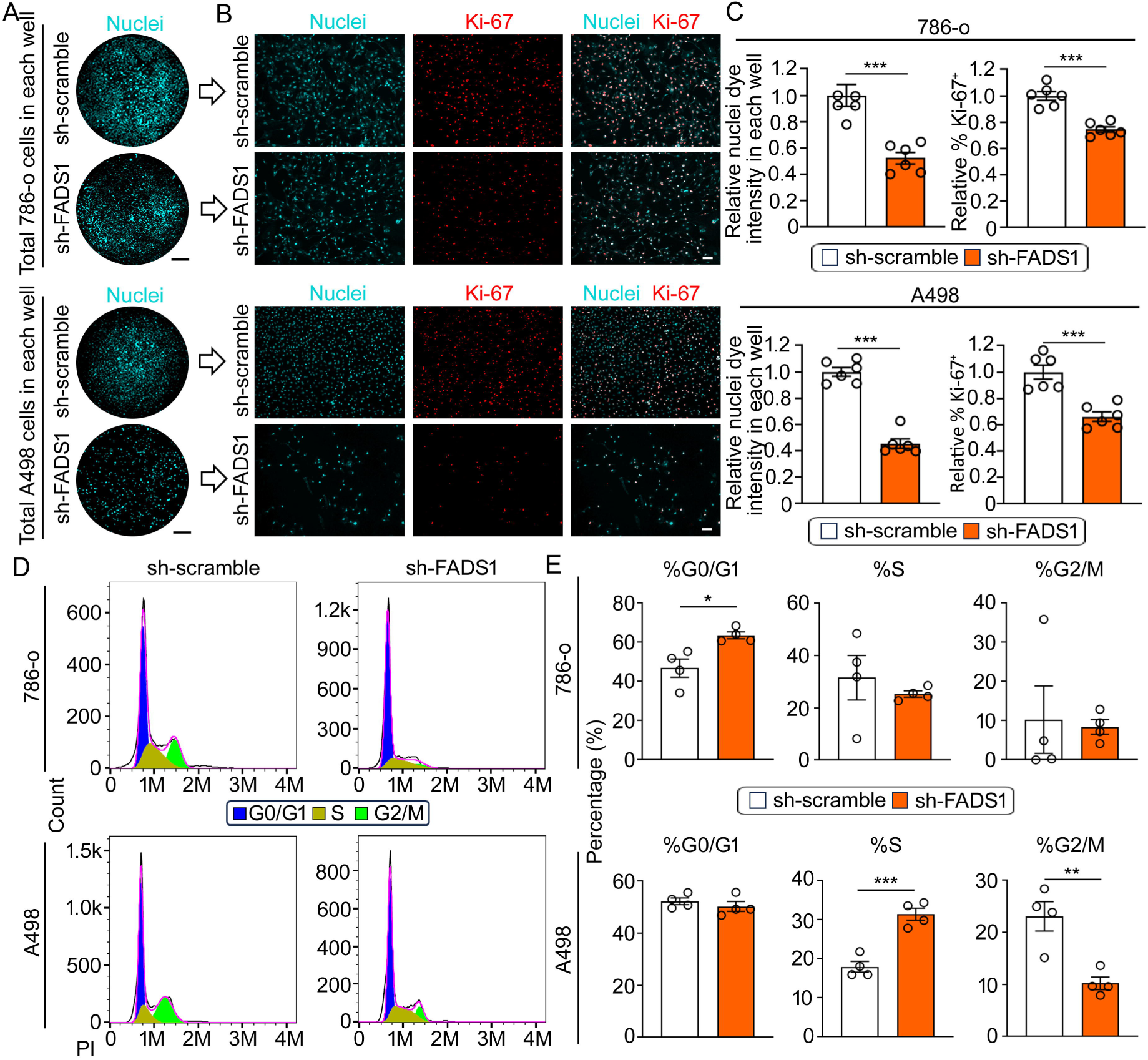
Suppression of *Fads1* gene inhibited human renal cancer cell growth. (A) Representative overview images of nuclei staining depict *Fads1* knockdown (sh-FADS1; bottom) and control (sh-scramble; top) 786-o (top) and A498 (bottom) cells in each well. Scale bar: 10 mm. (B) Representative immunofluorescence images showing expression of Ki-67 in sh-scramble and sh-FADS1 786-o (top) or A498 (bottom) cells. Scale bar: 100 μm. (C) The column bar graphs showing the relative quantification of whole nuclei staining intensity and the percentage of Ki-67 positive cells in sh-scramble and sh-FADS1 786-o or A498 cells (mean ± standard error). Data were normalized to the sh-scramble group. Statistical analysis performed using two-tailed unpaired Student’s t test. ***P<0.001. (D) Representative histogram showing the expression of the propidium iodide (PI) in sh-scramble and sh-FADS1 786-o (top) or A498 (bottom) cells. The different stages of cell cycle (G0/G1, S, G2/M stage) were determined by the expression of PI, as shown in the plot. (E) The column bar graphs showing the relative quantification of the percentage of each cell cycle stage (G0/G1, S, and G2/M) in sh-scramble and sh-FADS1 786-o (top) or A498 (bottom) cells (mean ± standard error). Data were normalized to the sh-scramble group. Statistical analysis conducted using two-tailed unpaired Student’s t test. *P<0.05; **P<0.01; ***P<0.001.

### Inhibition of FADS1 induces ER stress and ATF3 expression in renal cancer cells

Previous studies have demonstrated that lipid desaturation plays an important role in ER homeostasis. Inhibiting the desaturation process of saturated fatty acids leads to ER stress (16–19). Thus, we hypothesized that inhibition of LC-PUFA desaturation by targeting FADS1 would also induce ER stress and subsequently hinder the cancer cell growth. To test this hypothesis, we examined the expression of ER stress-related genes (*ATF3, ATF4, ATF6, CHOP, BIP, IRE1*, and *XBP1*) in cells treated with a FADS1 inhibitor. Our results showed that in FADS1 inhibitor-treated 786-o cells (**Fig. 3A & 3B**), all ER stress-related genes exhibited a significant increase when compared to the control group. FADS1 inhibitor-treated A498 cells exhibits a mixed result, with only ATF3 exhibiting a statistically significant increase (**Fig. 3C & 3D**). However, when assessing the expression of ER stress-related proteins in 786-o and A498 cells with or without FADS1 inhibitor treatment, we observed significant increase in protein expression of both ATF3 and ATF4 in FADS1 inhibitor-treated 786-o cells (**Fig. 3E & 3F**). FADS1 inhibitor-treated A498 cells also exhibit a significantly increased protein expression of ATF3 as well as a 1.6-fold increase in ATF4 protein, though not statistically significant. Interestingly, FADS1 inhibitor-treated A498 cells also show a significant increase in ATF6 protein level. (**Fig. 3G & 3H**). Taken together, inhibition of FADS1 in renal cancer cells promoted ER stress, with the likelihood that the ATF family of genes, especially ATF3, plays a key role in mediating this effect.

**Figure 3.**
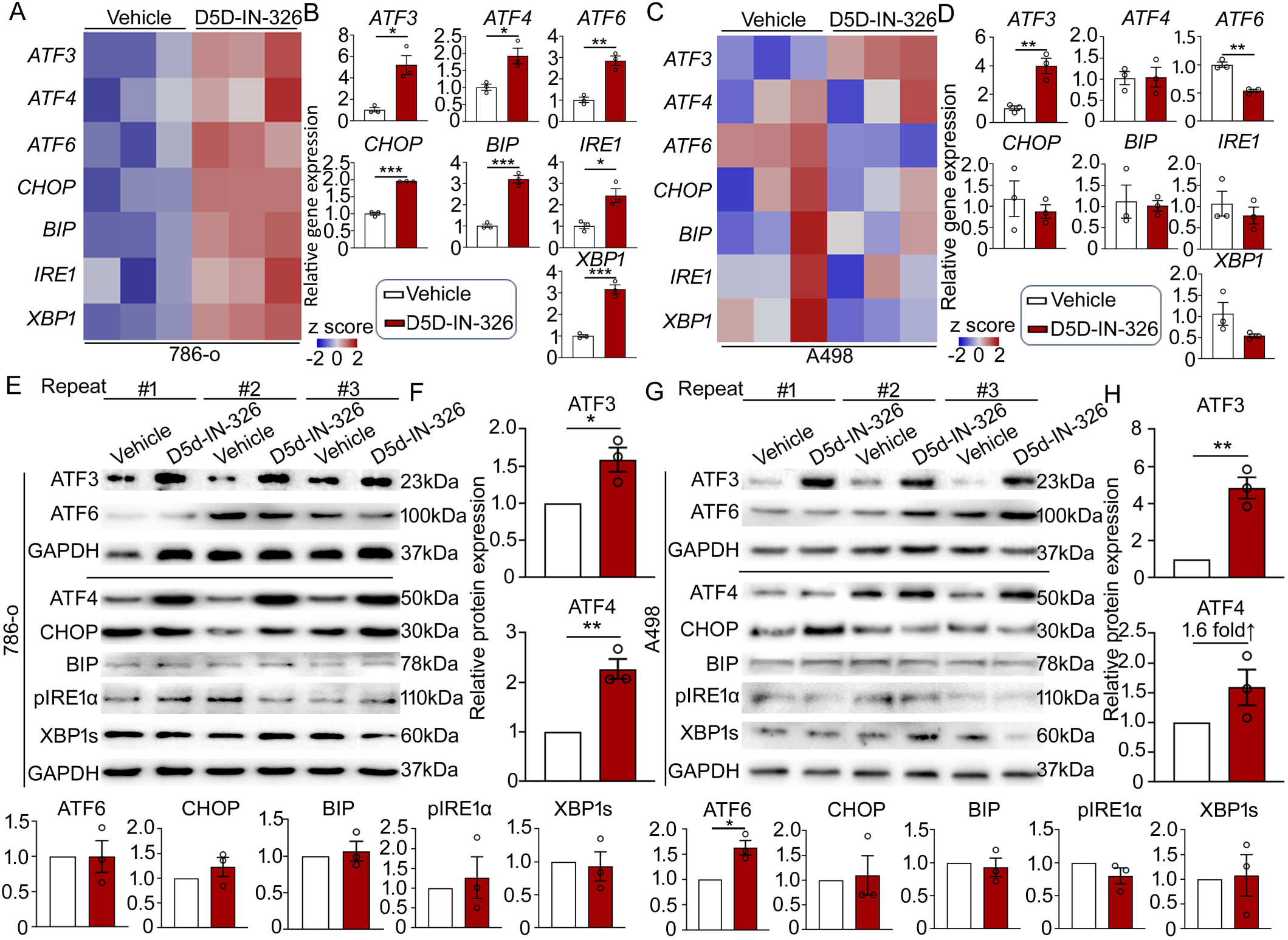
Inhibition of FADS1-triggered ER Stress activation in human renal cancer cells. (A) Heatmaps and (B) the column bar graphs showing the relative gene expression levels of ER stress-associated genes (*ATF3*, *ATF4*, *ATF6*, *CHOP*, *BIP*, *IRE1,* and *XBP1*) in 786-o cells or A498 cells (C & D) treated with 2,000 nM D5D-IN-326 (vehicle as control) for 48 hours (mean ± standard error). Data were normalized to the vehicle group. Statistical analysis performed using two-tailed unpaired Student’s t test. *P<0.05; **P<0.01; ***P<0.001. (E-H) Western blot images illustrating the expression of ER stress-associated proteins (ATF3, ATF6, ATF4, CHOP, BIP, pIRE1α, and XBP1s; GAPDH as housekeeping protein) in 786-o (E) and A498 (G) cells treated with 2,000 nM D5D-IN-326 (vehicle as control) for 48 hours. Showing here are results of assays independently repeated for three times. The column bar graphs showing the relative quantification of ER stress-associated proteins in 786-o (F) and A498 (H) cells treated with 2,000 nM D5D-IN-326 (vehicle as control) for 48 hours (mean ± standard error). Data were normalized to the vehicle group. Statistical analysis performed using two-tailed unpaired Student’s t test. *P<0.05; **P<0.01.

### FADS1 plays critical roles(s) in protecting renal cancer cells from extreme ER stress

In order to further explore the relationship between FADS1 and ER stress, we treated cells with two ER stress inducers, tunicamycin (TM) and thapsigargin (TG), to trigger ER stress response in wild type 786-o and A498 cells. As expected, after 18 hrs of treatment with TM or TG, our data show that ER stress markers start to be activated, though the pattern of such an induction in the two cell lines is different. However, FADS1 expression remains unchanged. After 24 hours of treatment with TM or TG, FADS1 transcription is significantly activated in both cells treated with TM or TG, and more ER stress markers are further induced. Again, among all ER stress markers ATF3 is consistently induced by TM and TG in both cell lines. (**Fig. 4A, 4B, 4C, & 4D**). These findings suggest that *FADS1* expression is induced by ER stress in renal cancer cells.

**Figure 4.**
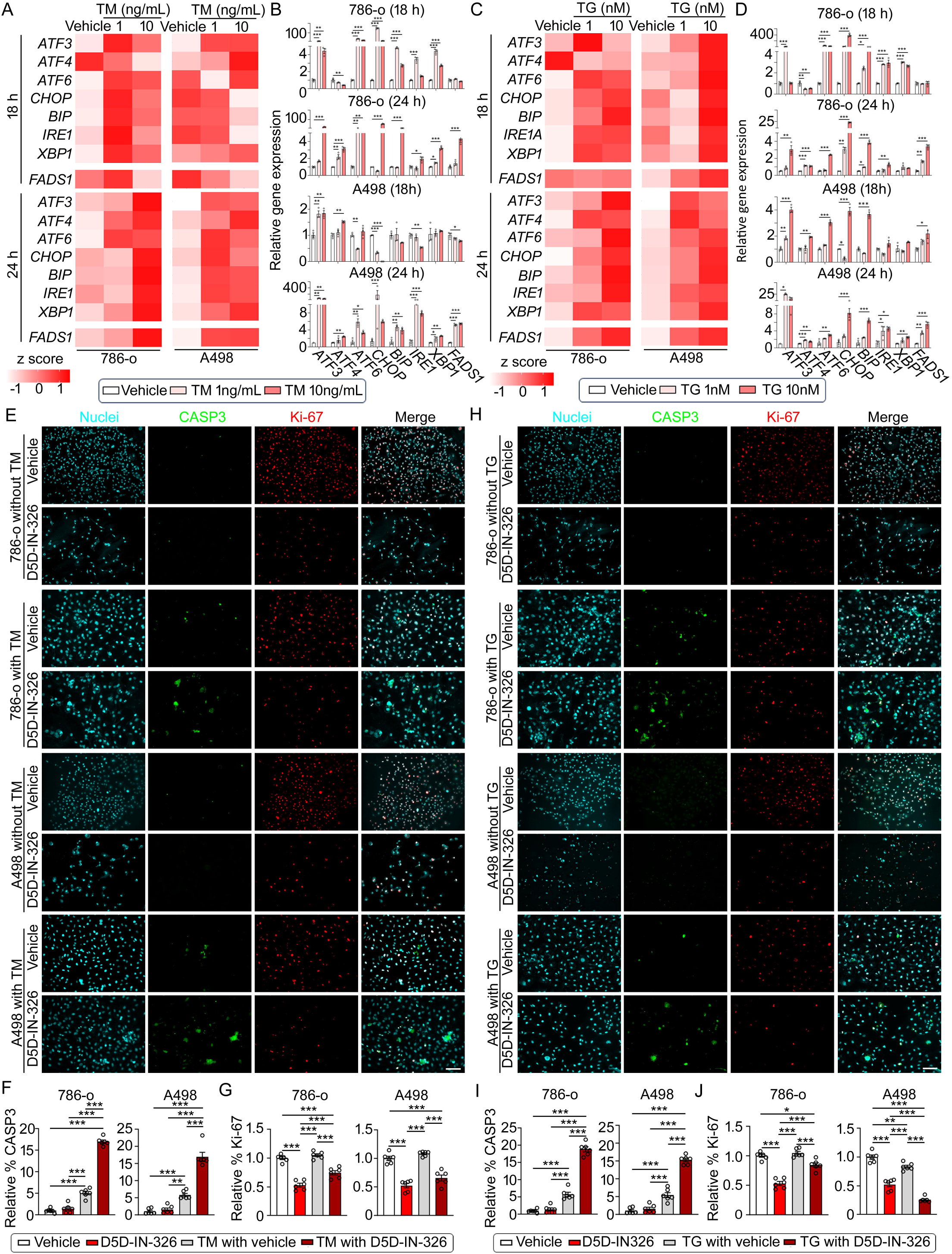
Increasing expression of *FADS1* reversed ER-stress induced human renal cancer cell apoptosis and slowed growth. (A&C) Heatmaps and (B&D) the column bar graphs illustrating the relative gene expression levels of ER stress-associated genes (*ATF3*, *ATF4*, *ATF6*, *CHOP*, *BIP*, *IRE1,* and *XBP1*) and *FADS1* in 786-o (left, A,C; top, B,D) and A498 (right, A,C; bottom, B,D) cells treated with gradient concentrations of Tunicamycin (TM) (vehicle, 1 ng/mL, and 10 ng/mL) (A&B) or Thapsigargin (TG) (vehicle, 1 nM, and 10 nM) for 18 hours and 24 hours (mean ± standard error). Data were normalized to the vehicle group. Statistical analysis conducted using Tukey’s multiple comparisons test. *P<0.05; **P<0.01; ***P<0.001. (E-J) Representative immunofluorescence images displaying expression of CASP3 and Ki-67 in 786-o (top images) and A498 (bottom images) cells treated with 2,000 nM D5D-IN-326 (vehicle as control) for 48 hours, with or without 18-hour pre-treatment with 1 ng/mL TM (E) or 1 nM TG (H). Scale bar: 100 μm. Bar graphs showing the relative quantification of the percentage of CASP3-positive cells (F) and Ki-67 positive cells (G) human renal cancer cells (786-o & A498) with or without TM or TG (i&j) pretreatment subjected to D5D-IN-326 treatment (vehicle as control) for 48 hours (mean ± standard error). Data were normalized to the vehicle group. Statistical analysis conducted using Tukey’s multiple comparisons test. **P<0.01; ***P<0.001.

Cancer cells often experience various extrinsic and intrinsic perturbations which lead to ER stress (33,34). Mild ER stress typically results in activation of signaling pathways that rescue cancer cells from stressful conditions. However, extreme or persistent ER stress may be unresolvable, especially when different cells possess different sensitivities to ER stress, which can lead to cell cycle arrest and apoptosis. (35–37). To determine if FADS1 is critical for protection of cancer cells from acute ER stress and the maintenance of cancer cell survival and growth, we pretreated 786-o and A498 cells with TM (1 ng/ml) or TG (1 nM) for 18 h. Subsequently, cells were co-treated with each of the ER stress inducers and the FADS1 inhibitor (2,000 nM) for additional 48 h. The co-treatment resulted in a significant increase in cell apoptosis, characterized by the elevation of CASP3-positive cells in TM and TG-treated 786-o and A498 cells compared to cells with TM/TG or D5D-IN-326 treatment alone (**Fig. 4E, 4F, 4H, & 4I**). Furthermore, FADS1 inhibition led to decreased number of Ki-67+ cells, though co-treatment with TM or TG does not show a significant synergistic effect (except for the TG and D5D-IN-326 co-treated A498 cells), possibly due to that the ER stress inducers *per se* do not exhibit impact on cell proliferation during the timeframe. (**Fig. 4E, 4G, 4H, & 4J**). These findings suggest that FADS1 function is critical for maintaining ER homeostasis following the induction of ER stress, and inhibiting FADS1 sensitizes cellular response to ER stress, inducing cell apoptosis.

### FADS1 regulation of renal cancer cell proliferation depends on ATF3

Given the significant increase in ATF3 expression in response to FADS1 inhibition, we set out to further investigate the potential role of ATF3 in mediating the impact of FADS1 inhibition on tumor cell growth. For this purpose, ATF3-specific shRNA to knock down ATF3 in 786-o and A498 cells were used. After optimizing ATF3 knockdown in these models (**Supplementary Fig. S2 & S3**), cell growth and proliferation following FADS1 inhibitor treatment was examined. Total cell number and the percentage of Ki-67^+^ cells were significantly lower in FADS1 inhibitor-treated 786-o and A498 cells compared to untreated control. However, upon ATF3 knockdown (ATF3-KD or sh-ATF3), proliferation was restored with the treatment of FADS1 inhibitor (**Fig. 5A, 5B, 5C, & 5D**). We further conducted cell cycle arrest assays to determine if ATF3 played a role in FADS1 inhibition-induced cell cycle arrest. Flow cytometry analysis indicated that FADS1 inhibition-induced cell cycle arrest was rescued, at least in part, by ATF3 gene knockdown (**Fig. 5E & 5F**). To investigate whether ATF3 knockdown could mitigate FADS1 inhibition-induced ER stress, we evaluated the expression of ER stress-related genes. FADS1 inhibition resulted in increased ER stress gene expression, which was significantly reduced by ATF3 gene knockdown in both cell lines (**Fig. 5G, 5H, 5I & 5J**). Taken together, our findings suggest that ATF3 plays a critical role in mediating the effect of FADS1 inhibition on ER stress, cell cycle and proliferation.

**Figure 5.**
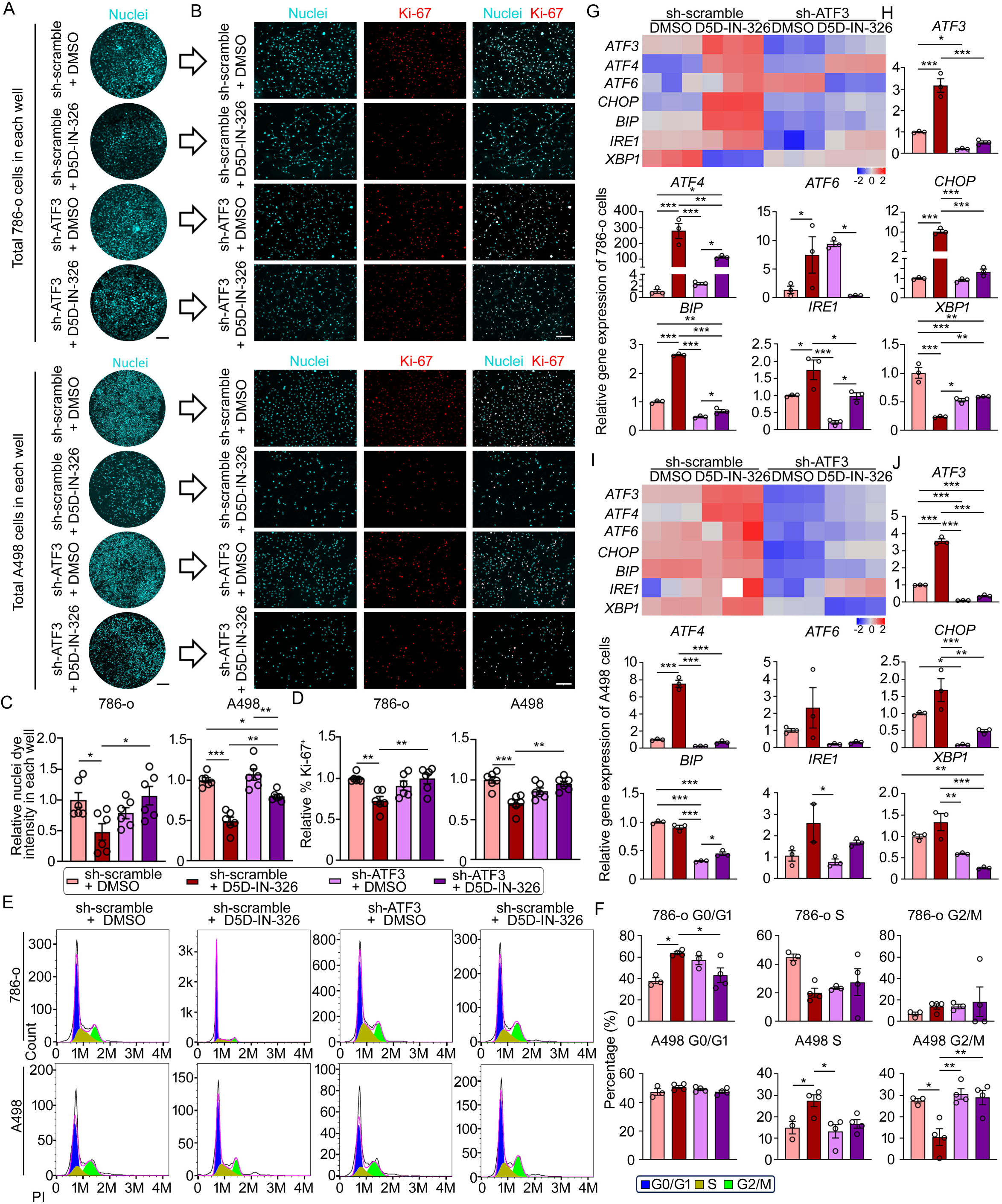
ATF3 mediates the reduction in human renal cancer cell proliferation upon FADS1 inhibition. (A) Representative images of nuclei staining showing *Atf3* knockdown (sh-ATF3) in 786-o (top) or A498 (bottom) cells treated with D5D-IN-326 (sh-scramble or sh-ATF3 with vehicle or D5D-IN-326) in each well. Scale bar: 10 mm. (B) Representative immunofluorescence images illustrating expression of Ki-67 in cells treated with sh-scramble with vehicle, sh-scramble with D5D-IN-326, sh-ATF3 with vehicle, and sh-ATF3 with D5D-IN-326 in 786-o (top) or A498 (bottom) cells. Scale bar: 100 μm. (C) Relative quantification of whole nuclei staining intensity and (D) the percentage of Ki-67 positive cells (mean ± standard error). Data were normalized to the sh-scramble+DMSO group. Statistical analysis conducted using Tukey’s multiple comparisons test. *P<0.05; **P<0.01; ***P<0.001. (E) Representative histogram showing the expression of the propidium iodide (PI) in sh-scramble and sh-FADS1 treated 786-o (top graphs) or A498 (bottom graphs) cells with or without D5D-IN-326 treatment. The different stages of cell cycle (G0/G1, S, G2/M stage) were determined by the expression of PI, as shown in the plot. (F) The column bar graphs showing the quantification of the percentage of each cell cycle stages (G0/G1, S, and G2/M) in cells treated with sh-scramble with vehicle, sh-scramble with D5D-IN-326, sh-ATF3 with vehicle, and sh-ATF3 with D5D-IN-326 of 786-o (top graphs) or A498 (bottom graphs) cells (mean ± standard error). Statistical analysis conducted using one-way ANOVA followed by Tukey’s multiple comparisons test. *P<0.05; **P<0.01. (G) Heatmaps and (H) column bar graphs depicting relative gene expression levels of ER stress-associated genes (*ATF3*, *ATF4*, *ATF6*, *CHOP*, *BIP*, *IRE1,* and *XBP1*) in cells treated with sh-scramble with vehicle, sh-scramble with D5D-IN-326, sh-ATF3 with vehicle, and sh-ATF3 with D5D-IN-326 786-o and (I & J) A498 cells (mean ± standard error). Data were normalized to the vehicle group. Statistical analysis conducted using Tukey’s multiple comparisons test. *P<0.05; **P<0.01; ***P<0.001.

To further verify the role of ATF3 in mediating the FADS1-KD-induced cell proliferation and cell cycle arrest, we introduced both ATF3- and FADS1-specific shRNA into 786-o and A498 cells. In both cells, sh-FADS1 and sh-ATF3 knockdown significantly reduced FADS1 and ATF3 mRNA and protein expression, respectively. Cells co-treated with sh-FADS1+sh-ATF3 significantly reduced mRNA and protein expression of both ATF3 and FADS1 (**Supplementary Fig. S2 & S3**). Subsequently, we assessed cell growth and proliferation in these genetically edited cells. Consistent with our previous findings, the total cell number and the percentage of Ki-67+ cells in FADS1-KD 786-o and A498 cells were significantly lower relative to the scramble control groups. However, these effects were significantly reversed in cells with both FADS1 and ATF3-knockdown (**Fig. 6A-F**). Similarly, we also demonstrated that ATF3 knockdown partly rescued the cell cycle arrest induced by FADS1-KD (**Fig. 6g & 6h**). These results further demonstrate that ATF3 is a key factor mediating the effects of FADS1 inhibition in cell proliferation.

**Figure 6.**
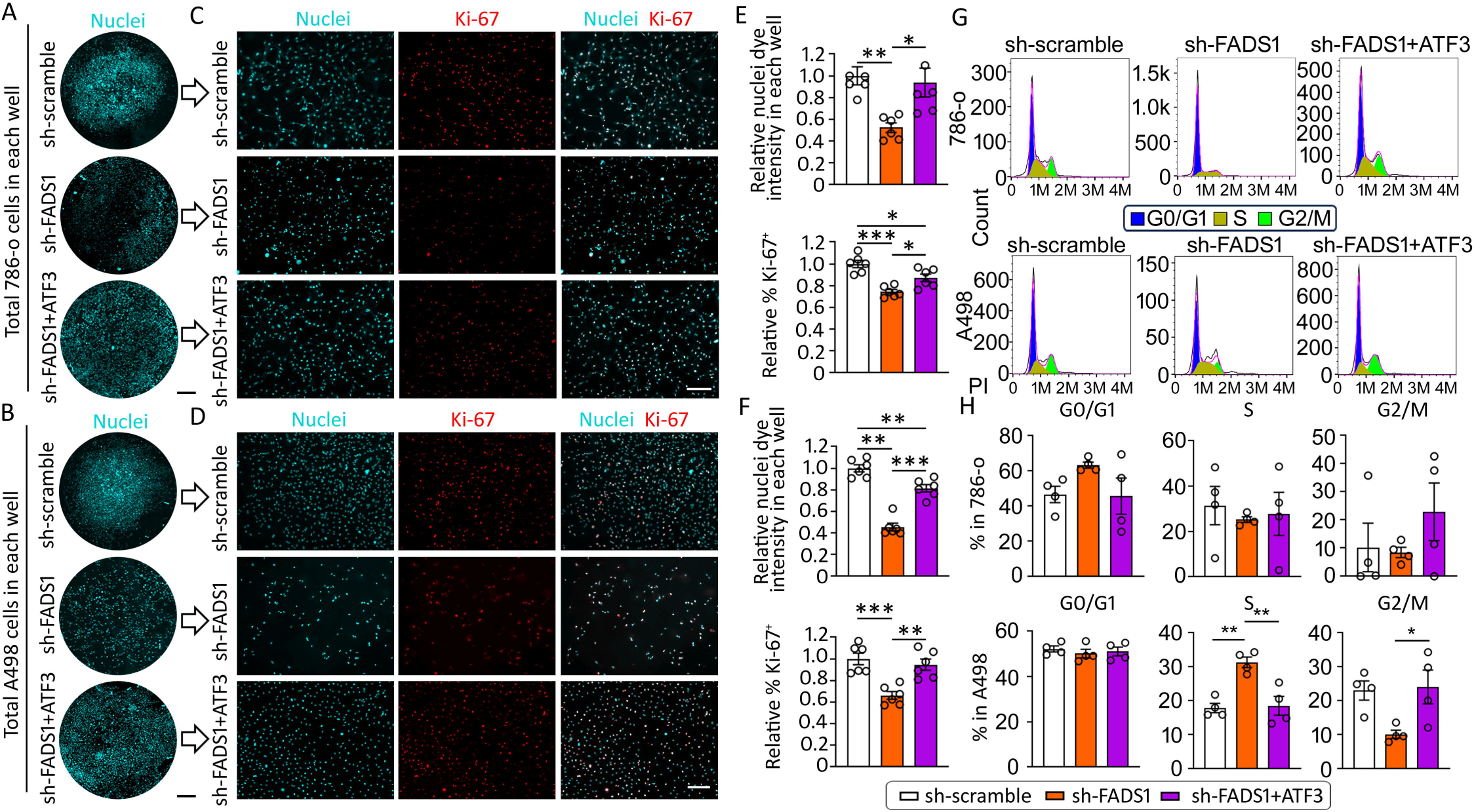
Co-knockdown of *ATF3* and *FADS1* rescues reduced cell proliferation induced by *Fads1* inhibition. (A) Representative overview images of nuclei staining showing scramble control (sh-scramble), *Fads1* knockdown (sh-FADS1), and *ATF3* & *FADS1* combination knockdown (sh-ATF3 & FADS1) of 786-o cells or (B) A498 cells in each well. (C) Representative immunofluorescence images illustrating expression of Ki-67 in cells treated with sh-scramble, sh-FADS1, and sh-ATF3 & FADS1 786-o cells or (D) A498 cells. Scale bar: 10 mm in (A) and 100 μm in (B). (E) The column bar graphs showing the relative quantification of whole nuclei staining intensity and percentage of Ki-67 positive cells in sh-scramble, sh-FADS1, and sh-ATF3 & FADS1 786-o cells or (F) A498 cells (mean ± standard error). Data were normalized to the sh-scramble group. Statistical analysis was conducted using Tukey’s multiple comparisons test. *P<0.05; **P<0.01; ***P<0.001. (G) Representative histogram showing the expression of the propidium iodide (PI) in sh-scramble, sh-FADS1, and sh-ATF3 & FADS1 786-o (top) or A498 (bottom) cells. The different stages of cell cycle (G0/G1, S, G2/M stage) were determined by the expression of PI, as shown in the plot. (H) The column bar graphs showing the percentage of each cell cycle stage (G0/G1, S, and G2/M) in sh-scramble, sh-FADS1, and sh-ATF3 & FADS1 786-o (top) or A498 (bottom) cells (mean ± standard error). Statistical analysis conducted using Tukey’s multiple comparisons test. *P<0.05; **P<0.01.

### FADS1 inhibition decreases biosynthesis of nucleotides and UDP-N-Acetylglucosamine

To further understand the potential metabolic remodeling that FADS1 inhibition may cause, we conducted comprehensive metabolomic analyses using 786-o sh-scramble and 786-o sh-FADS1 cells. As a proof-of-concept for establishing the role of FADS1 in lipid metabolism, we noted that FADS1 knockdown significantly impacted pathways involving fatty acid metabolism, including biosynthesis, elongation, and degradation, characterized by significantly reduced levels of acetyl-CoA and CoA. Interestingly, pathways regulating amino sugar and nucleotide sugar metabolism were also enriched (**Supplementary Fig. S5A**). Notably, FADS1 knockdown significantly reduced the level of nucleotides and their precursors, e.g., dATP, dGTP, dTDP, AICAR, etc. More importantly, FADS1 knockdown significantly downregulated the level of UPD-N-Acetylglucosamine (UDP-GlcNAc), a critical substrate of protein glycation in the ER (**Supplementary Fig. S5B**). Reduced UDP-GlcNAc has been shown to induce protein unfolding or misfolding, activating the unfolded protein response, a key step in ER stress homeostasis (38). These findings suggest that FADS1 inhibition exerts a pronounced impact on nucleotide biosynthesis and protein folding, which may account for the cell cycle arrest and ER stress observed in our experiments.

### FADS1 knockdown inhibits tumor growth in vivo

To assess the impact of FADS1 on tumor development *in vivo*, we injected wild-type (WT) and FADS1-KD 786-o cells (**Supplementary Fig. S6A**) into the flank of male and female severe combined immunodeficient (SCID) mice to generate tumors (**Fig. 7A**). We monitored tumorigenesis over a course of 55 d. The size of FADS1-KD tumors was significantly smaller than those formed by wild-type cells, both in female and male mice, with more significant difference in female mice (**Fig. 7B**). After 55 d, tumors were collected for further analyses (**Fig. 7C; Supplementary Fig. S6B**). The weight of tumor xenografts derived from the FADS1-KD or control cells demonstrated a similar trend as the tumor size (**Fig. 7D & 7E**). Taken together, these results demonstrate that reduced FADS1 function limits tumor formation *in vivo*, suggesting inhibiting FADS1 might have a therapeutic potential for cancer.

**Figure 7.**
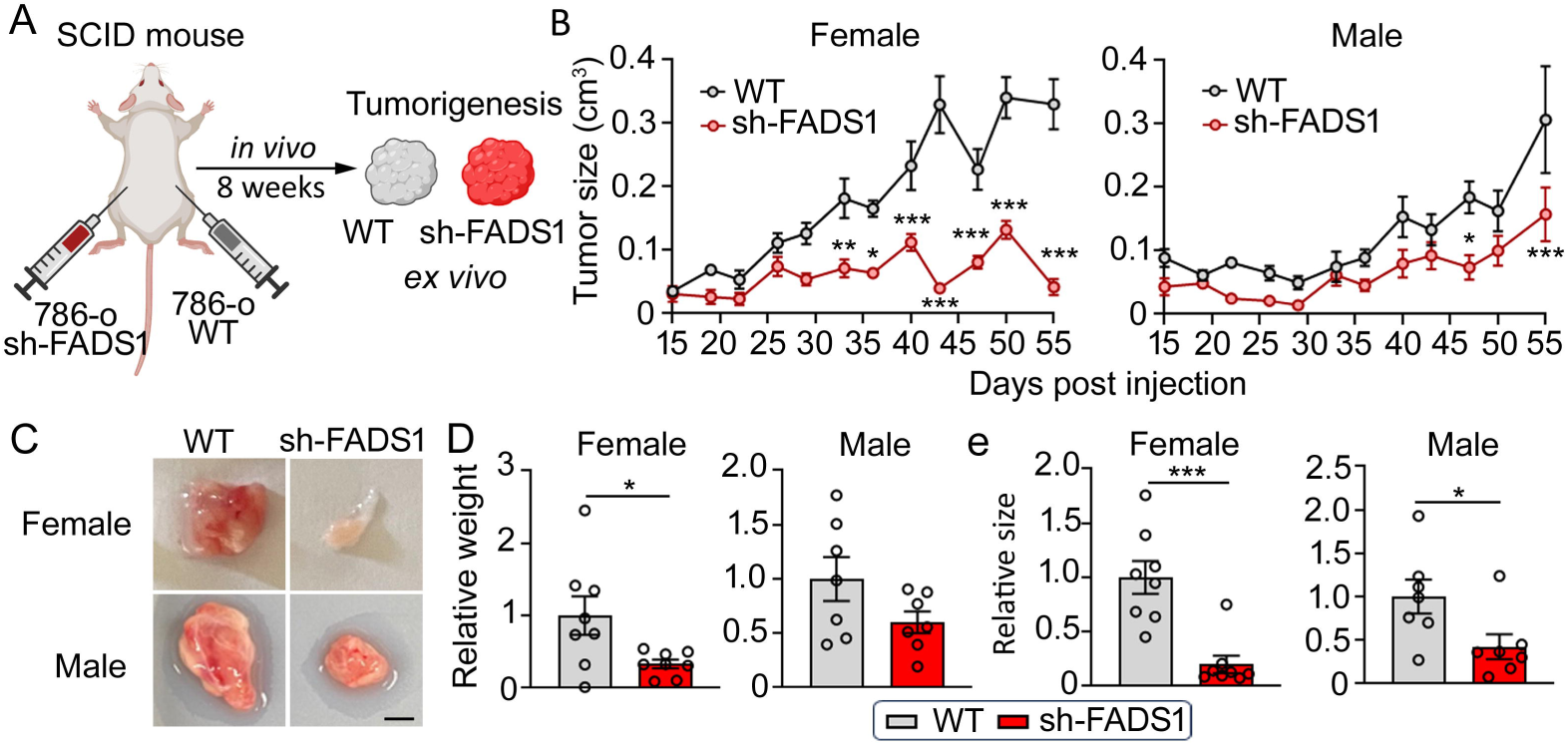
FADS1 Knockdown Reduces Renal Cancer Cell Growth *In Vivo*. (A) Experimental diagram illustrating the human renal cancer cell xenograft study. FADS1 knockdown (sh-FADS1) 786-o cells were injected into the left flank of SCID mice, while wild type (WT) 786-o cells were injected into the right flank. Tumor growth was monitored for 8 weeks. (B) The graphs depicting average tumor size (cm^3^) of sh-FADS1 786-o cells and WT 786-o cells in female and male severe combined immunodeficiency disease (SCID) mice up to 55 days post injection (mean ± standard error). Statistical analysis performed using two-tailed unpaired Student’s t test. *P<0.05; **P<0.01; ***P<0.001. (C) Representative images of extracted subcutaneous tumors from male and female mice. Scale bar: 0.25 cm. (D) Relative tumor weight (grams) and (E) size (cm^3^) in WT and KD subcutaneous tumors post extraction (mean ± standard error). Data were normalized to the WT group. Statistical analysis performed using two-tailed unpaired Student’s t test. *P<0.05; ***P<0.001.

## Discussion

Increasing evidence suggests that FADS1 activity and PUFA metabolism are closely linked to cancer initiation, progression and metastasis (29). Our previous research has demonstrated that increased FADS1 gene expression is significantly correlated with reduced cancer patient survival among multiple types of cancers, and especially in kidney cancer patients (13). We also found that FADS1 mRNA levels were significantly increased in RCC tumors compared to normal tissues. The FADS1 transcription level is even higher in metastatic and recurrent tumors (13), highlighting its potential role in cancer biology and disease progression. These early observations prompted us to further explore the causal role of FADS1 in RCC biology as well as the potential underlying mechanism. In this study, we demonstrate that FADS1 knockdown or pharmacological inhibition suppresses cell proliferation, alters cell cycle in RCC cells, and reduces tumor formation *in vivo*. We further provide evidence that FADS1 is essential for supporting cancer cell growth by protecting cancer cells from persistent ER stress, which is mediated by a key transcription factor ATF3. Our data provide new knowledge regarding the essential role of FADS1 in RCC and demonstrates the therapeutic potential of targeting FADS1 for renal cancer.

We found that FADS1 inhibition-induced suppression of cell proliferation is mediated by ER stress. The ER plays a pivotal role as a central organelle responsible for protein synthesis, folding, and maturation (39). When cells are under stress, misfolded or unfolded proteins accumulate, which triggers the unfolded protein response (UPR) and initiates ER stress (39). Cancer cells are often exposed to stressful microenvironmental conditions e.g. hypoxia and nutrient deprivation, as well as intrinsic conditions such as rapid metabolism and overproduction of reactive oxygen species (ROS), which typically lead to the activation of ER stress (40–42). ER stress responsive signaling may activate downstream pathways to support tumor growth, metastasis, and adaptation to anti-cancer treatments (33,34,43). However, extreme or persistent conditions may induce unresolvable ER stress, leading to cell death. Logically, different cancer cells may exhibit different levels of tolerability to ER stress. There is a reciprocal relationship between lipids metabolism and ER homeostasis. While ER stress can disturb lipid synthesis, metabolism and intracellular trafficking, lipid supplementation and intracellular metabolism can also alter ER function and related signaling (44–46). Previous studies have demonstrated the important role of fatty acid desaturation in maintaining ER homeostasis and function. Increased lipids saturation, especially long-chain fatty acids saturation, induces ER stress (47–51), which has been attributed to a reduction in ER membrane fluidity (52), ER Ca(2+) depletion (53,54), overload of misfolded proteins (48), and direct interaction between lipids and ER signal proteins (50). Notably, limiting MUFA biosynthesis by blocking SCD1 has also been shown to activate ER stress and cell death in ccRCC and other cancers (55–58). FADS1 is a rate-limiting enzyme responsible for delta-5 desaturation of both omega-3 and omega-6 LC-PUFAs, with its immediate metabolic products being AA and EPA. Our study demonstrated for the first time that the inhibition of FADS1 leads to a reduced conversion of DGLA to AA, and induces an ER stress response. Reciprocally, chemically induced ER stress also increases the transcription of FADS1, while inhibition of FADS1 reduces cells tolerance to ER stress inducers. These data suggest that FADS1 expression is required to rescue RCC cells from persistent ER stress. This is noteworthy, because kidney cancers are known to be sensitive to hypoxia signals as well as anti-angiogenic agents, suggesting a potentially high basal level of ER stress in these cancers. Our previous analysis has shown an increased FADS1 expression in kidney cancers with a pattern of expression level in normal tissues<primary tumors<metastatic tumors<recurrent tumors (13). Combined with the observations from our current study, these data may reflect an escalating dependence of FADS1 expression in response to an increased ER stress level during the progression of kidney cancers. We have previously demonstrated that FADS1 expression is significantly associated with poor patient survival among all three independent cohorts of kidney cancer patients in TCGA (13). Our studies hence collectively identify FADS1 as a key gene supporting the growth of renal cancer cells and as a therapeutic target for cancer treatment.

Within the ER membrane, three primary pathways are known to become active during ER stress: PERK, ATF6, and IRE1 (59). Activation of these sensors sends signals to the nucleus, ultimately modifying cellular function and growth. Our data suggest that FADS1 inhibition predominantly activates the ATF4-ATF3 branch, which is under PERK-mediated UPR pathway, possibly culminating in increased ATF3 expression within the nucleus. ATF3, being a key transcription factor, exerts diverse roles in cellular metabolism and function, including regulation of glucose metabolism, lipogenesis, immune response, and modulation of ER stress-induced responses (60,61). In the context of cancer cells, ATF3 was demonstrated to have differential roles across various cancer types. For example, ATF3 functions as an oncogene in prostate cancer, where its high expression is associated with increased cell proliferation in response to androgen stimulation (61,62). In breast cancer, elevated ATF3 expression induces the expression of MMP13, TWIST, Slug, and Snail, thereby modulating tumor metastasis (61,63). In contrast, ATF3 acts as a tumor suppressor in lung cancer by inducing cancer cell apoptosis via activation of DR5 (61,64,65). Similar to lung cancer, ATF3 functions as a tumor suppressor in colon and liver cancer by limiting cell proliferation and inducing apoptosis (61,66,67). Our study supports ATF3 as a tumor suppressor in RCC that restrains renal cancer cell proliferation. How ATF3 regulates cell cycle and cell proliferation by mediating the effect of FADS1 inhibition remains to be further investigated. Interestingly, inhibiting SCD1 to block MUFA production was also shown to induce ER stress (68,69), in which ATF3 plays a critical role in mediating the downstream cellular responses in both cancer and other health conditions (15,58,70–72). Therefore, the findings in our present study suggest that LC-PUFA production, in part regulated by FADS1, plays a similar role to that of MUFA production. Collectively, our study indicates that fatty acids desaturation for either MUFA or PUFA production is a fundamental mechanism for maintaining ER homeostasis and function.

It should be noted that fatty acids may also directly contribute to cell proliferation and regulation of cell metabolism. In stark contrast to normal cells, cancer cells exhibit an insatiable demand for energy, membrane constituents, and regulatory factors to facilitate their rapid proliferation (73–75). Consequently, extensive metabolic reprogramming becomes imperative to sustain this aberrant growth. A plethora of studies have underscored the critical involvement of fatty acid metabolism, encompassing both biosynthesis and desaturation of fatty acids, in fueling cancer cell proliferation (76–79). More importantly, LC-PUFAs and their lipid derivatives e.g. eicosanoids, prostaglandins, and leukotrienes, have been broadly demonstrated to directly regulate cancer cell proliferation and cell cycle (7,80). Our study demonstrates that FADS1 inhibition or knockdown leads to reduced production of AA metabolites, which possibly further reduces the cellular levels of downstream AA derivatives. This potential lipidomic remodeling and its role in regulating cell proliferation and cell cycle cannot be disregarded and should be further explored in future studies.

We also noticed that FADS1 inhibition induces cell cycle arrest instead of cell death. This may be due to RCC cells, in culture, having a lower intrinsic baseline ER stress as compared to an RCC tumor *in vivo*. Therefore, cultured RCC cells might be less dependent on FADS1 expression. Indeed, when pre-treating the cells with ER stress inducers, FADS1 expression is increased, and FADS1 inhibition at this point induces cell apoptosis, as indicated by the increased CASP3+ cells. Also, some studies have demonstrated that FADS1 inhibition or reduced expression enhances cancer cell sensitivity (81). *In vivo*, due to the harsh microenvironment of kidney cancers, it is highly likely that cancer cells possess an elevated ER stress level, thus an escalated dependence on FADS1 function, as indicated by the aforementioned increased FADS1 expression in more advanced stages of RCC. Therefore, RCC tumors should be sensitive to FADS1 inhibition. Studies are ongoing in our lab to explore the potential impact of hypoxia, nutrient deprivation and existing anti-RCC drugs on ER stress in RCC cells, as well as the sensitivity of cells under these conditions to FADS1 inhibition.

To further investigate the causal factors mediating the impact of FADS1 inhibition on cell cycle arrest and ER stress, we conducted a metabolomic analysis in control and FADS1-KD 786-o cells. We observed a significant reduction in the levels of nucleotides and their related intermediate molecules in FADS1-KD cells. Previous studies have demonstrated that the availability of nucleotides is a critical factor in cell cycle progression (82–84). Therefore, the reduced nucleotides biosynthesis may restrain DNA synthesis, which explains the cell cycle arrest to G1-S phase following FADS1 inhibition. However, why FADS1-KD results in depletion of nucleotides requires further investigation. In addition to the impact on DNA synthesis, our study also demonstrates that FADS1-KD cells exhibit a decrease in the level of UDP-GlcNAc. This sugar nucleotide is the product of the hexosamine biosynthetic pathway (HBP) and is the key substrate of protein N-glycosylation in the ER lumen (85). Protein N-glycosylation is known to regulate protein folding and a diminished intracellular UDP-GlcNAc level may trigger the UPR signaling and induces ER stress (86). This finding underscores the intricate connections between FADS1 function, metabolic remodeling, and cell stress. Future experiments are needed to explore how FADS1 and ATF3 lead to the metabolic remodeling for these pathways.

In summary, our findings demonstrate that FADS1 inhibition effectively curtails renal cancer cell proliferation *in vitro* as well as tumor formation *in vivo*, primarily through the activation of the ATF3 axis-mediated ER stress. Our study highlights FADS1-ER stress as a key therapeutic target for renal cancer, paving the way for novel drug discovery and development.

## Acknowledgements

We acknowledge Dr. Jesscia Back and Eric VanBuren at Microscopy, Imaging, and Cytometry Resources Core of Wayne State University for their technical expertise and assistance with the flow cytometry assays and data analysis.

## Author contributions

Conceptualization: G.H., Z.L., I.P. W.L. Methodology: G.H., Z.L., K.Z., J.L., R.C., I.P., W.L. Validation: G.H., Z.L., W.L. Formal analysis: G.H., Z.L., W.L. Investigation: G.H., Z.L., M.H., A.W., Y.F., Y.J.,N.V., L.T., Z.P., W.L. Data curation: G.H., Z.L., M.H., A.W., Y.F., Y.J., N.V., L.T., Z.P., W.L. Writing-original draft: G.H., Z.L., W.L. Writing-review & editing: all authors. Project administration: W.L.

## Availability of data and materials

All data produced or examined within this study are comprehensively incorporated in both the manuscript and its supplementary documents.

## Declarations

### Ethics approval and consent to participate

All experiments involving mice were performed in accordance with the protocol (ID# IACUC-21-12-4269) approved by the institutional Animal Investigational Committee of Wayne State University and NIH guidelines.

### Consent for publication

All authors have agreed to publish this manuscript.

