## Supplemental figures and tables for "Targeting Fatty Acid Desaturase I Inhibits Renal Cancer Growth Via ATF3-mediated ER Stress Response"

**Supplemental Data**

**Supplemental Figure S1**

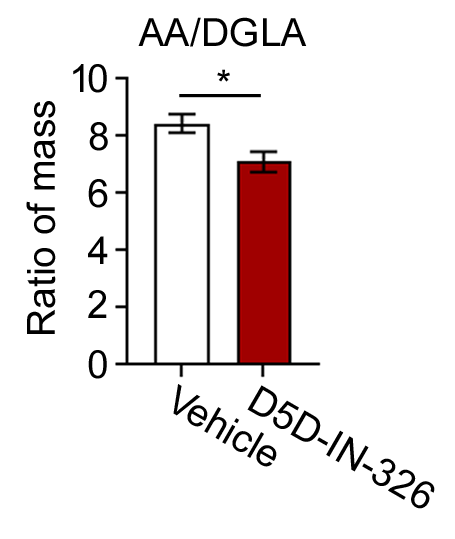

**Supplementary Figure S1. D5D-IN-326 inhibits Polyunsaturated Fatty Acid Desaturation.**

The column graph showing the ratio of AA/DLGA in 786-o cells with 2,000 nM D5D-IN326 treatment (vehicle as control) (mean ± standard error). Statistical analysis performed using two-tailed unpaired Student’s t test. *P<0.05.

**Supplemental Figure S2**

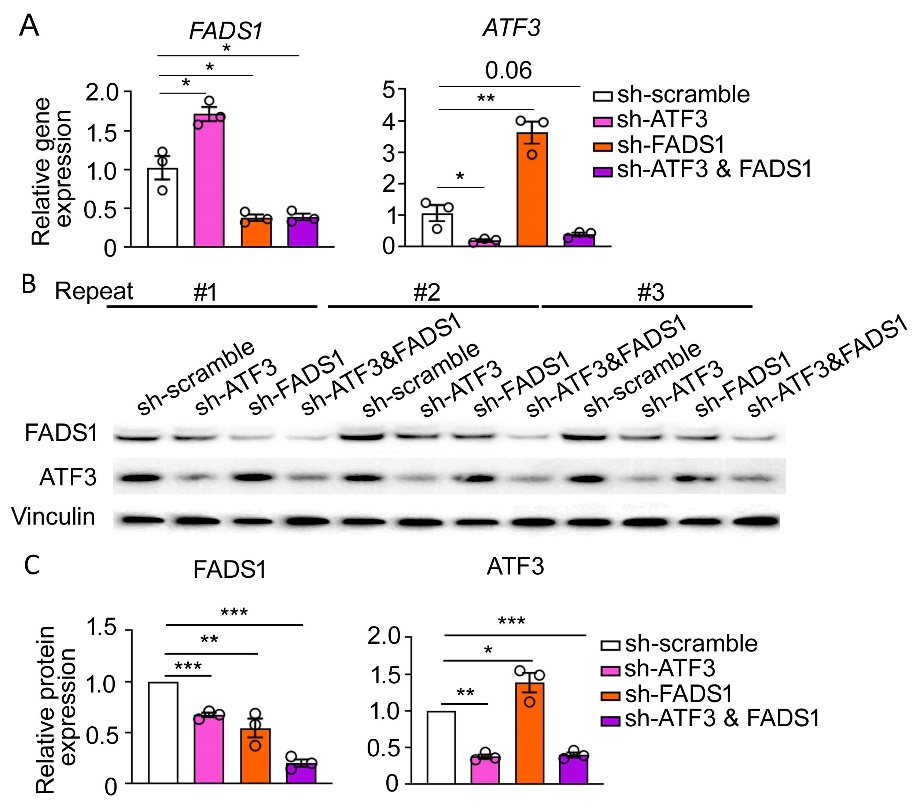

**Supplementary Figure S2. Evaluation of the sh-ATF3, sh-FADS1, and sh-ATF3 & FADS1 786-o cells**

(A) The column graph showing relative *ATF3* and *FADS1* gene expression in sh-scramble, sh-ATF3, sh-FADS1, sh-ATF3 & FADS1 786-o cells. Data were normalized to the sh-scramble group. Statistical analysis was conducted using two-tailed unpaired Student’s t test. *P<0.05; **P<0.01. (B) Western blot images illustrating expression of ATF3 and FADS1 protein (Vinculin as the housekeeping protein) in sh-scramble, sh-ATF3, sh-FADS1, sh-ATF3 & FADS1 786-o cells. (C) The column bar graphs showing the relative quantification of ATF3 and FADS1 proteins in sh-scramble, sh-ATF3, sh-FADS1, sh-ATF3 & FADS1 786-o cells based on Western blot analysis (mean ± standard error). Data were normalized to the sh-scramble group. Statistical analysis conducted using Statistical analysis conducted using two-tailed unpaired Student’s t test. *P<0.05; **P<0.01; ***P<0.001.

**Supplemental Figure S3**

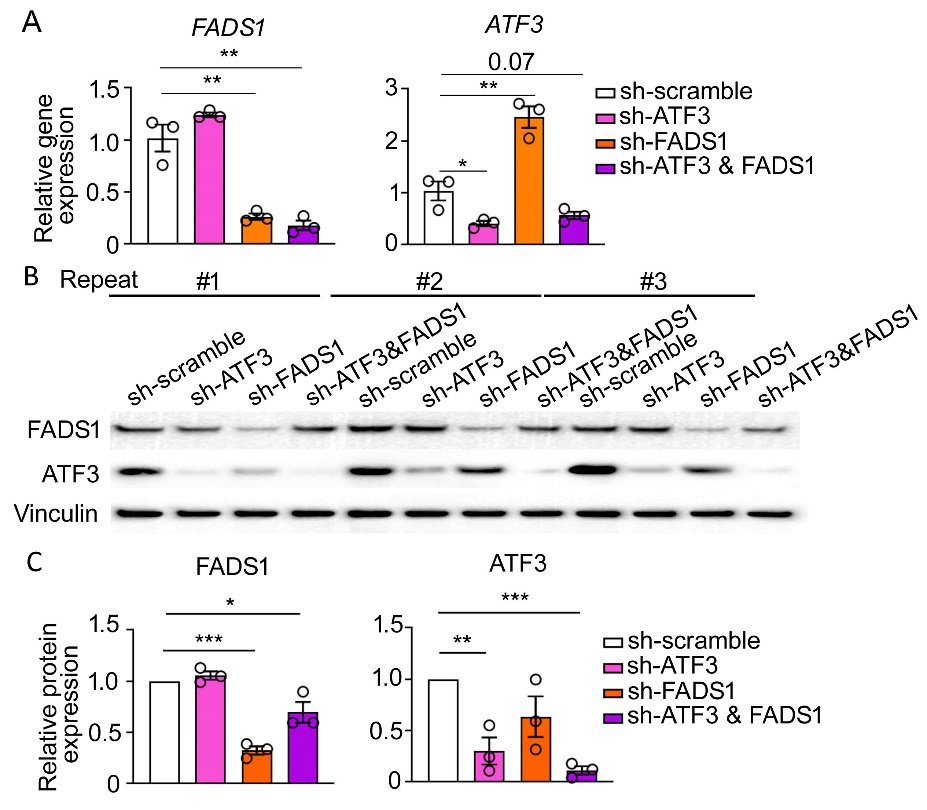

**Supplementary Figure S3. Evaluation of the sh-ATF3, sh-FADS1, and sh-ATF3 & FADS1 A498 cells**

(A) The column graph showing relative *ATF3* and *FADS1* gene expression in sh-scramble, sh-ATF3, sh-FADS1, sh-ATF3 & FADS1 A498 cells. Data were normalized to the sh-scramble group. Statistical analysis was conducted using two-tailed unpaired Student’s t test. *P<0.05; **P<0.01. (B) Western blot images illustrating expression of ATF3 and FADS1 protein (Vinculin as the housekeeping protein) in sh-scramble, sh-ATF3, sh-FADS1, sh-ATF3 & FADS1 A498 cells. (C) The column bar graphs showing the relative quantification of ATF3 and FADS1 proteins in sh-scramble, sh-ATF3, sh-FADS1, sh-ATF3 & FADS1 A498 cells based on Western blot analysis (mean ± standard error). Data were normalized to the sh-scramble group. Statistical analysis conducted using Statistical analysis conducted using two-tailed unpaired Student’s t test. *P<0.05; **P<0.01; ***P<0.001.

**Supplemental Figure S4**

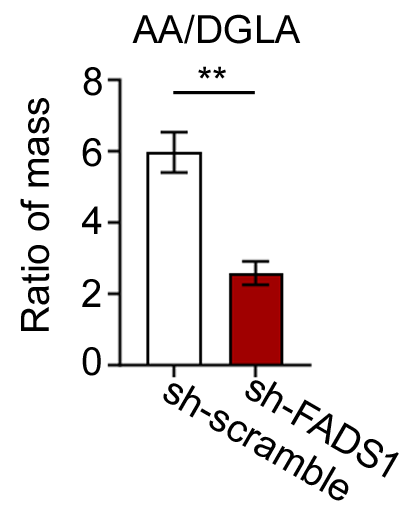

**Supplementary Figure S4. FADS1 knockdown Inhibits Polyunsaturated Fatty Acid Desaturation**

The column graph showing the ratio of AA/DLGA in sh-scramble and sh-FADS1 786-o cells (mean ± standard error). Statistical analysis performed using two-tailed unpaired Student’s t test. **P<0.01.

**Supplemental Figure S5**

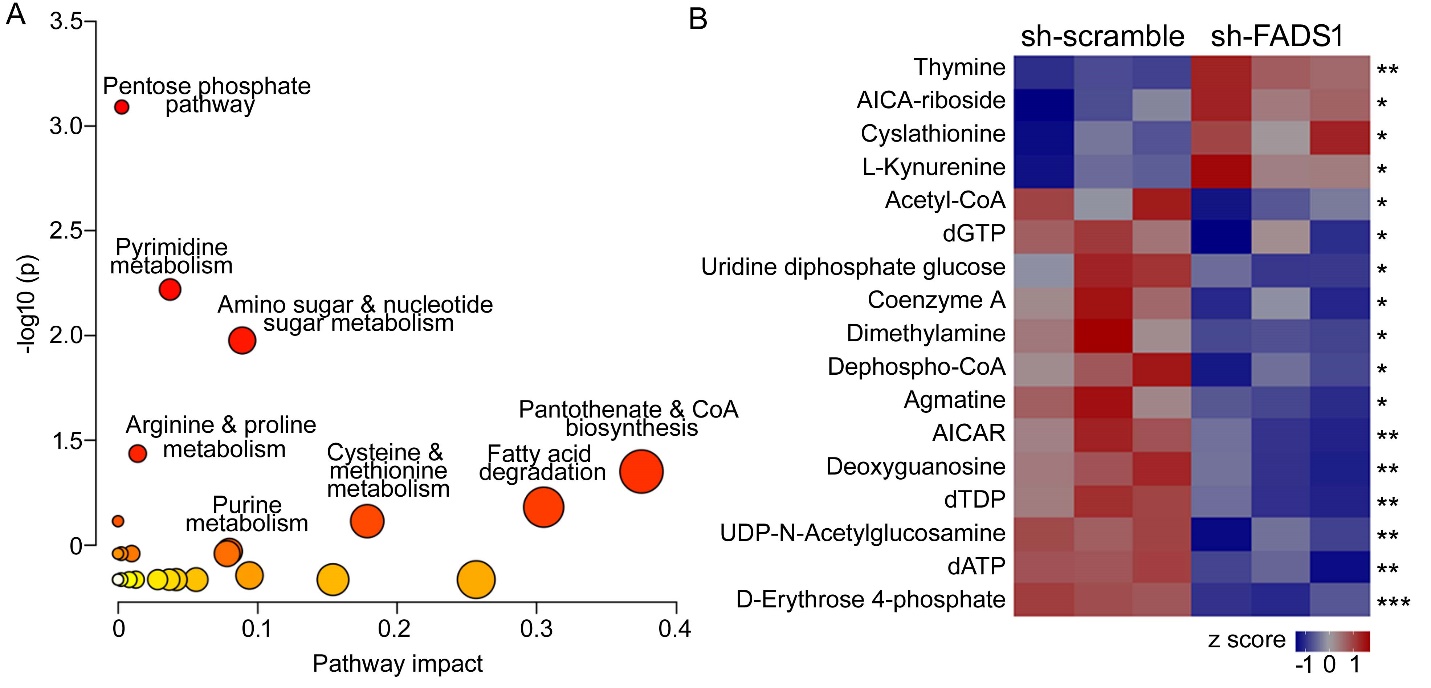

**Supplementary Figure S5: Impact of FADS1 knockdown on metabolomic profile of 786-o cells.**

(A) Metabolomic pathway analysis plot revealing enriched (p<0.05) metabolic pathways in 786-o cells with FADS1 knockdown. (B) Heatmap plot illustrating the metabolites with significant change in their levels in between sh-scramble and sh-FADS1 786-o cells (relative to z score). Statistical analysis was conducted using a two-tailed unpaired Student’s t-test. *P<0.05, **P<0.01, ***P<0.001.

**Supplemental Figure S6**

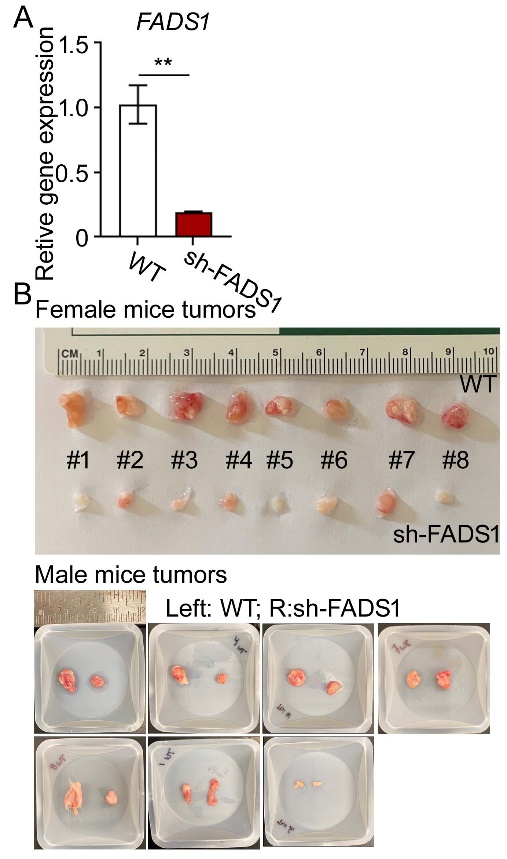

**Supplementary Figure S6: Images of extracted tumors from male and female mice**

(A) The column graph showing the *FADS1* gene expression in WT and sh-FADS1 786-o cells (mean ± standard error; *in vivo* study only). Data were normalized to the WT group. Statistical analysis performed using two-tailed unpaired Student’s t test. **P<0.01. (B) The images of extracted subcutaneous tumors from (top) female (8 pairs of tumors; upper tumors were 786-o WT and lower tumors were 786-o sh-FADS1) and (bottom) male mice (7 pairs of tumors; one mouse did not develop any tumor; left tumors were 786-o WT and right tumors were 786-o sh-FADS1).

Supplemental Table S1. Sequences for the shRNA experiments (All sequences are provided in 5’ to 3’ orientation)

| **Name** | **Target** | **Hairpin sequence** |
| --- | --- | --- |
| sh-ATF3  (SCBT; Cata# sc-29757-V) | *ATF3* | GATCCCGACGAGAAAGAAATAAGATTCAAGAGATCTTATTTCTTTCTCGTCGTTTTT |
|  |  | GATCCGTGTGAATGCTGAACTGAATTCAAGAGATTCAGTTCAGCATTCACACTTTTT |
|  |  | GATCCCTCTCCACTCAATGTCTTATTCAAGAGATAAGACATTGAGTGGAGAGTTTTT |
| sh-FADS1  (SCBT; Cata# sc-96474-V) | *FADS1* | GATCCGGATAGGTATGACCTATGTTTCAAGAGAACATAGGTCATACCTATCCTTTTT |
|  |  | GATCCCTCTAGGCATATTGATCATTTCAAGAGAATGATCAATATGCCTAGAGTTTTT |
|  |  | GATCCCCATGGAGAGGTTTGTCAATTCAAGAGATTGACAAACCTCTCCATGGTTTTT |
| sh-FADS1  (*in vivo*) | *FADS1* | GATCCCGCCTTGTGAAGAAGTATATGAATTCAAGAGA TTCATATACTTCTTCACAAGG TTTTT |

Supplemental Table S2. Primers for the RT-qPCR analyses

| **Gene** | **NCBI ID** | **Forward primer** | **Reverse primer** |
| --- | --- | --- | --- |
| *FADS1* | 3992 | CCTGGAAAGCAACTGGTTTGTG | GAAGGCAGACTTGTGGACATTG |
| *PPIA* | 5478 | AGGTCCCAAAGACAGCAGAA | GAAGTCACCACCCTGACACA |
| *ATF3* | 467 | CCTCTGCGCTGGAATCAGTC | TTCTTTCTCGTCGCCTCTTTTT |
| *ATF4* | 468 | CCCTTCACCTTCTTACAACCTC | TGCCCAGCTCTAAACTAAAGGA |
| *ATF6* | 22926 | AGCAGCACCCAAGACTCAAAC | GCATAAGCGTTGGTACTGTCTGA |
| *CHOP* | 1649 | AGGAACCAGGAAACGGAAACAGA | TCTCCTTCATGCGCTGCTT |
| *BIP* | 3309 | CATCACGCCGTCCTATGTCG | CGTCAAAGACCGTGTTCTCG |
| *IRE1α* | 2081 | GCCGAAGTTCAGATGGAATC | ATCTGCAAAGGCCGATGA |
| *XBP1* | 7494 | GCAGGTGCAGGCCCAGTTGT | TGGGTCCAAGTTGTCCAGAAT |

Supplemental Table S3. Primary antibodies for immunofluorescence assays

| **Antibody** | **Species** | **Dilution** | **Company (Catalog #)** |
| --- | --- | --- | --- |
| ATF3 | Mouse | 1:100 | Santa Cruz Biotechnology (sc-518032) |
| CASP3 | Rabbit | 1:200 | Abcam (ab32351) |
| FADS1 | Rabbit | 1:200 | Abcam (ab126706) |
| Ki-67 | Rat | 1:100 | Thermofisher (14-5698-82) |

Supplemental Table S4. Secondary antibodies for immunofluorescence assays

| **Antibody** | **Conjugate** | **Dilution** | **Company (Catalog #)** |
| --- | --- | --- | --- |
| Donkey anti mouse | Alexa 647 | 1:500 | Jackson ImmunoResearch (715-606-150) |
| Donkey anti rabbit | Alexa 488 | 1:500 | Jackson ImmunoResearch (711-546-152) |
| Donkey anti rat | Cyanine Cy3 | 1:500 | Jackson ImmunoResearch (712-166-150) |

Supplemental Table S5. Primary antibodies for western blot assay

| **Antibody** | **Species** | **Dilution** | **Company (Catalog #)** |
| --- | --- | --- | --- |
| ATF3 | Rabbit | 1:500 | Abcam (ab254268) |
| ATF4 | Rabbit | 1:1000 | Abcam (ab270980) |
| ATF6 | Mouse | 1:1000 | Novus biologicals (NBP1-40256) |
| BIP | Rabbit | 1:1000 | Novus biologicals (NBP1-06277) |
| CHOP | Rabbit | 1:1000 | Novus biologicals (NBP2-13172) |
| FADS1 | Rabbit | 1:1000 | Abcam (ab126706) |
| GAPDH | Rabbit | 1:1000 | Cell signaling (2118s) |
| pIRE1α | Rabbit | 1:1000 | Novus biologicals (NB100-2323) |
| XBP1 | Rabbit | 1:1000 | Novus biologicals (NBP1-77681) |

Supplemental Table S6. Secondary antibodies for western blot assays

| **Antibody** | **Dilution** | **Company (Catalog #)** |
| --- | --- | --- |
| Anti-mouse, HRP linked | 1:1000 | Cell signaling (7076s) |
| Anti-rabbit, HRP linked | 1:1000 | Cell signaling (7074s) |
